# Cellular-level phenotyping of tumor-immune microenvironment (TiME) in patients in vivo reveals distinct inflammation and endothelial anergy signatures

**DOI:** 10.1101/2021.06.10.447835

**Authors:** Aditi Sahu, Teguru Tembo, Kivanc Kose, Anthony Santella, Anabel Alfonso, Madison Li, Miguel Cordova, Melissa Gill, Christi Fox, Salvador Gonzalez, Amber Weiching Wang, Nicholas Kurtansky, Pratik Chandrani, Piyush Kumar, Shen Yin, Haaris Jilani, Paras Mehta, Cristian Navarrete-Dechent, Gary Peterson, Kimeil King, Stephen Dusza, Ning Yang, Shuaitong Li, William Phillips, Anthony Rossi, Allan Halpern, Liang Deng, Melissa Pulitzer, Ashfaq Marghoob, Chih-Shan Jason Chen, Milind Rajadhyaksha

**Author notes:** Corresponding authors: Milind Rajadhyaksha, Aditi Sahu, Dermatology Service, Memorial Sloan Kettering Cancer Center, 530E, 74^th^ street, New York-10021.

## Abstract

Immunotherapies have shown unprecedented clinical benefits in several malignancies^1–3^. However, clinical responses remain variable and unpredictable, indicating the need to develop predictive platforms that can improve patient stratification^4^. Phenotyping of tumors into hot, altered, or cold^5^ based on assessment of only T-cell infiltration in static tumor biopsies provides suboptimal prediction of immunotherapy response^6,7^. *In vivo* dynamic mechanisms within the tumor microenvironment such as tumor angiogenesis and leukocyte trafficking^5,8,9^ also play a central role in modulating anti-tumor immunity and therefore immunotherapy response. Here, we report novel tumor immune microenvironment (TiME) phenotyping *in vivo* in patients with non-invasive spatially-resolved cellular-level imaging based on endogenous contrast. Investigating skin cancers as a model, with reflectance confocal microscopy (RCM) imaging^10^, we determined four major phenotypes with variable prevalence of vasculature (Vasc) and inflammation (Inf) features: Vasc^hi^Inf^hi^, Vasc^hi^Inf^lo^, Vasc^lo^Inf^hi^ and Vasc^med/hi^Inf^lo^. The Vasc^hi^Inf^hi^ phenotype correlates with high immune activation, exhaustion, and vascular signatures while Vasc^hi^Inf^lo^ with endothelial anergy and immune exclusion. Automated quantification of TiME features demonstrates moderate-high accuracy and correlation with corresponding gene expression. Prospectively analyzed response to topical immunotherapy show highest response in Vasc^lo^Inf^hi^, and reveals the added value of vascular features in predicting treatment response. Our novel *in vivo* cellular-level imaging and phenotyping approach can potentially advance our fundamental understanding of TiME, develop robust predictors for immunotherapy outcomes and identify novel targetable pathways in future.

## Main

Immunotherapy, especially immune checkpoint blockade therapy, has revolutionized cancer management by providing near durable responses in several cancers^1–3^. However, only a subset of patients derives long-term benefit, highlighting a clinical need to develop effective biomarkers for patient stratification^4,11,12^. Phenotyping of tumors into hot, cold or altered based on T-cell infiltration in tumor center and margin; PD-L1 expression; and tumor mutation burden are important determinants for immunotherapy in solid cancers^13,14^. Although hot versus cold tumor phenotyping has shown some association with response, not all inflamed phenotypes respond to treatment. Thus, immune-cell infiltration is necessary but insufficient for inducing anti-tumor immunity^7,15^. Clearly, tumors utilize additional mechanisms for evading immune response while establishing an immune-suppressive microenvironment, complicating patient stratification strategies because of dynamic tumor/host immune crosstalk and baseline tumor biology^13,16,17^.

Tumor vasculature (blood and lymphatic vessels) serve important immunomodulatory roles, and contribute to the immune evasion of tumors^18^. Angiogenesis promotes immune evasion through induction of a highly immunosuppressive TiME by inhibiting dendritic cell (DC) maturation, T-cell development and function, and most importantly, limiting effector immune cells access to tumors by modulating leukocyte trafficking^19^. In addition, tumor vasculature can display decreased expression of adhesion molecules, and non-responsiveness to inflammatory cytokines—attributes of vascular endothelial anergy. By downregulating trafficking of effector immune cells, endothelial anergy contributes to ineffective anti-tumor immune responses and immune evasion^20–22^.

Towards addressing the dynamic, complex and highly interdependent vascular-inflammation axis inside the TiME, *in vivo* phenotyping based on a combination of dynamic vascular and immune features, rather than *ex vivo* phenotyping based on static pathological evaluation of tumor infiltrating cells alone may facilitate a deeper understanding and achieve higher predictive power for patient stratification for immunotherapies. High-resolution non-invasive *in vivo* imaging is fundamental to this combination phenotyping, since static *ex vivo* analyses on patient tissue are limited in recapitulating dynamic vascular and the continuous, evolving cellular-level crosstalk between the immune system and cancer ^12,23^. We report novel phenotypes detected *in vivo* using reflectance confocal microscopic (RCM) imaging. RCM is a high-speed (pixel times ~ 0.10 μs, frame rates 10-30 per second) cellular-level label-free imaging approach based on backscattered light and endogenous tissue contrast^10,24^. Large image mosaics (64 mm^2^ in 50 s) imaged to a depth of ~0.25 mm enables spatial resolution of TiME features. RCM is routinely used for real-time skin cancer diagnosis and management at the bedside. Two studies have briefly reported RCM imaging of vessels and leukocyte trafficking in humans^25,26^. Using skin cancer as a model, we define TiME phenotypes using six features: vessel diameter, vessel density, leukocyte trafficking, intratumoral inflammation, peritumoral inflammation and perivascular inflammation and report their subsequent molecular correlation with inflammatory, angiogenic, trafficking and tumor-intrinsic signatures. In a prospective pilot study, we also investigate the relative importance of TiME features and phenotypes in predicting response to topical immunotherapy.

## Results

Patients with suspected skin cancers (n=27) who underwent biopsy for diagnosis or treatment were included for phenotyping. Gene expression was successfully performed in a subset of patients (n=14). In addition, a subset (n=13) of superficial basal cell carcinoma (BCCs) were prescribed topical immunotherapy (Imiquimod) and their response was correlated with TiME features and phenotype (Fig 1A). The agreement for manual evaluation of RCM TiME features (Fig S1, supplementary videos 1-3) by two independent readers was performed. The evaluation was also correlated with analogous histopathological features by a board-certified dermatopathologist (Table 1). Substantial to almost perfect agreement (k=0.62–1.0) was observed for most RCM features. Good to very good agreement (AC_1_: 0.74-1.0) was found between histopathology and average RCM evaluation. Unsupervised multivariate analysis of TiME features was performed using Principal Component Analysis (PCA) and clustering on PCA data revealed four major phenotypes: Vasc^hi^Inf^hi^, Vasc^hi^Inf^lo^, Vasc^lo^Inf^hi^ and Vasc^med/hi^Inf^lo^ (Fig 1B, Fig S2A). The *in vivo* phenotypes were correlated with the total area of CD3^+^ T cells (to assess with respect to T-cell based pathological phenotyping^5^) and area of tertiary lymphoid structures (TLS) (Fig S3), a hallmark of an inflamed micro-environment and positive cancer outcomes^27^ (Fig 1C).

**Figure 1.**
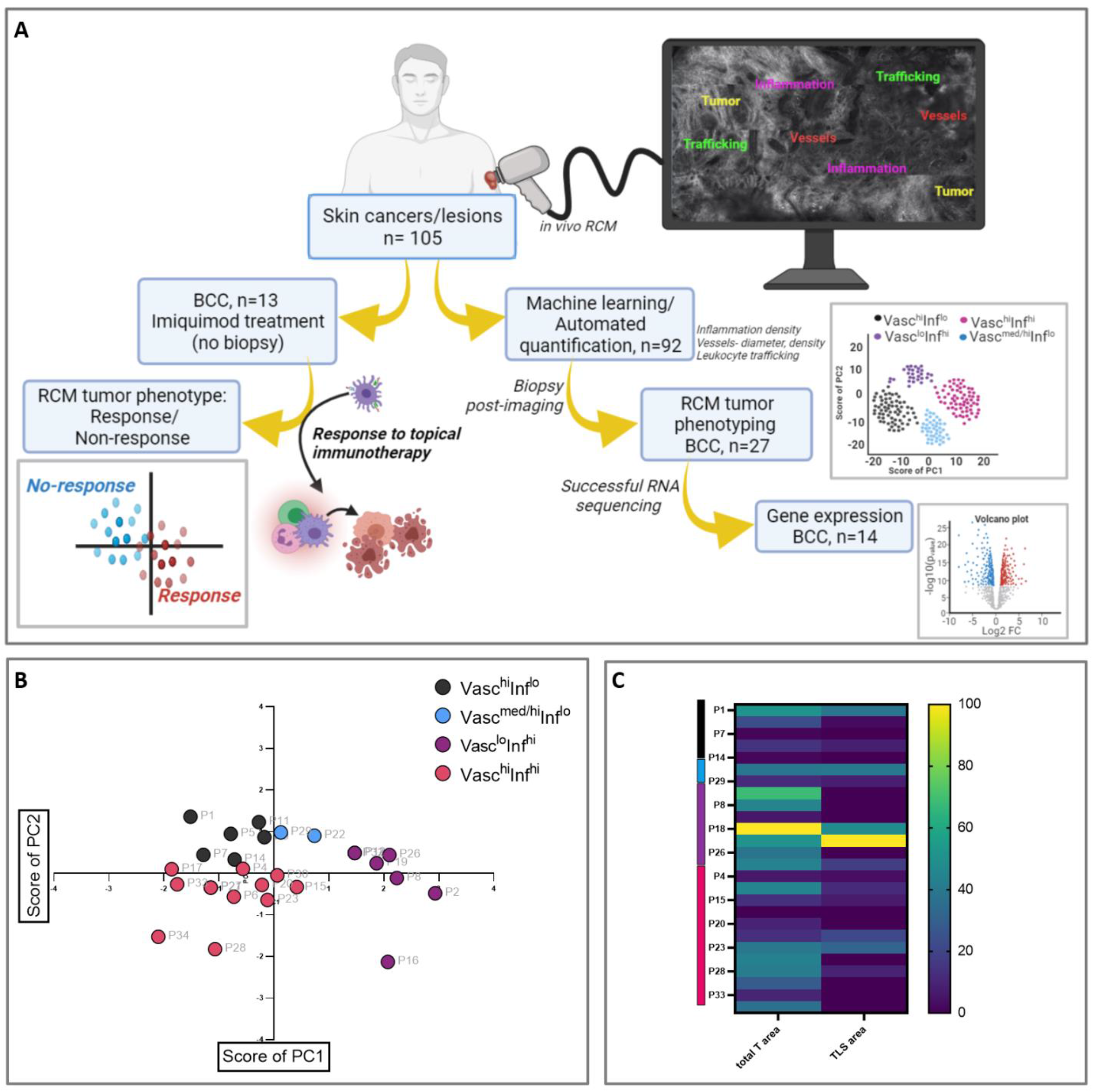
Overview of the research strategy, TiME phenotyping in patients followed by immune correlation. **A.** *In vivo* RCM imaging on patients with skin cancers or skin lesions visiting the Dermatology Service at MSKCC was performed to characterize tumor, immune and vascular features. Data was used for machine learning and automated quantification of inflammation density, vessel diameter and density, and frequency of leukocyte trafficking. RCM phenotyping using unsupervised analysis was done on a subset of patients who received a biopsy following imaging. IHC for CD3, CD20 was performed on these available biopsies and correlated with RCM phenotypes. Bulk RNA-sequencing after RNA extraction from FFPE tissues was also explored on these tissues, only 14 samples yielded sufficient quality and quantity for bulk RNA sequencing. The *in vivo* TiME phenotypes were correlated using DGEA, CIBERSORT and GSEA. In addition, selected BCC patients with confirmed RCM diagnoses of superficial subtype were selected for topical immunotherapy imiquimod (treatment regimen includes 5x/week for 6 weeks). The patients were imaged 6 months after end of treatment to confirm tumor clearance and classify patients as responders (complete tumor regression) and non-responders (incomplete or no tumor regression). The phenotypes and TiME features associated with response were identified. Created on www.Biorender.com **B.** Unsupervised PCA demonstrate four major phenotypes: Vasc^hi^Inf^hi^, Vasc^hi^Inf^lo^, Vasc^lo^Inf^hi^ and Vasc^med/hi^Inf^lo^. **C.** Heatmap of differences in CD3^+^ T cell and TLS area between the 4 phenotypes. BCC: basal cell carcinoma, PCA: principal component analysis, FFPE: formalin fixed paraffin embedded, DGEA: differential gene expression analysis, GSEA: gene set enrichment analysis and TLS: tertiary lymphoid structures.

**Table 1.** High agreement confirms the reproducibility and presence of TiME features on histopathology. Substantial to almost perfect agreement (k=0.62-1.0) was observed for RCM features between 2-readers, k for peritumor inflammation and perivascular inflammation agreement could not be computed due to 100% prevalence. Good to very good agreement (AC_1_: 0.74-1.0) was found in the binary analysis between average RCM evaluation and dermatopathologist grading of features.

| **TiME Feature** | **RCM 2-Reader Agreement**<br>**(Cohen's kappa)** | **Histology vs. Average 2-**<br>**reader (Binary)** |
| --- | --- | --- |
| Number of vessels | 0.62 | 0.84 |
| Dilated vessel | 0.72 | 0.89 |
| Trafficking | 0.76 | 0.72 |
| Intratumor inflammation | 1.00 | 1.00 |
| Peritumor inflammation | NA | 0.97 |
| Mucin | 0.93 | 0.74 |
| Perivascular inflammation | NA | 1.00 |

Transcriptomic analysis on samples with sufficient RNA quality and quantity for bulk RNA-sequencing (Fig S4A) enabled phenotype correlation with gene expression profiling. TiME phenotyping on this patient subset (n=14) were coalesced into two main groups (Inf^hi^ and Vasc^med/hi^) (Fig 2A) for group-level transcriptomic analysis. Phenotyping into 4 groups was also investigated (Fig S2B, C). Phenotyping on gene expression data was also explored for inflammation, angiogenesis^28^ and trafficking gene signatures. Similar tendency of classification, especially between the Inf^hi^ and Vasc^med/hi^ phenotypes was observed in inflammation (driven mainly by TNFAIP2 expression), angiogenesis (mainly SPARCL1) and trafficking (mainly CXCL12) (Fig S2E-F). Next, CIBERSORT^29^, differential gene expression analysis (DGEA) and pathway analysis, and gene set enrichment analysis (GSEA) were undertaken on Inf^hi^ and Vasc^med/hi^ phenotypes Unsupervised k-means clustering on CIBERSORT output (Fig 2B, Fig S4B) shows two distinct Inf^hi^ and Vasc^med/hi^ phenotypes similar to RCM (Fig 2A), with one misclassified patient in each cluster (P21 and P22). DGEA on the CIBERSORT output on the Inf^hi^ and Vasc^med/hi^ phenotypes found 11 significant differentially expressed genes (Fig S4C). Relative immune cell proportions indicated higher CD4^+^ memory resting, CD4^+^ memory active and M1, M2 macrophages in Inf^hi^ (Fig S4C). DGEA on the total transcriptome guided by the RCM TiME phenotypes (Inf^hi^ and Vasc^med/hi^) found 114 differentially expressed genes that separated RCM TiME phenotypes into the same 2 groups with hierarchical cluster analysis (HCA) (Fig S4E), 85 genes were overexpressed in the Inf^hi^ while 29 genes in the Vasc^med/hi^ cluster (Fig 2C). Pathway analysis using available gene sets demonstrate enrichment of mainly pro-inflammatory pathways in Inf^hi^ while enrichment of leukocyte-endothelial interactions, ephrin B2 pathway, vasodilation and neovascularization in the Vasc^med/hi^ (Fig 2D). Gene set enrichment analysis (GSEA) on the DGEA data show enrichment of immune and angiogenesis signatures in the Inf^hi^ cluster (Fig S4D). HCA on Nanostring Pancancer Immune Panel Genes^30^ (Fig S4F) indicates presence of four phenotypes of which P7, P14 and P17 (Vasc^med/hi^) were found to have comparatively lower expression of immune genes as compared to P5, P6, P21, P23, P36 (Inf^hi^). We also investigated phenotyping on 3 GEO datasets^31–33^ using PCA and DGEA towards analyzing similar phenotypic trends in additional diverse cohorts (Fig S5). Specific differences in major inflammation, vascular (angiogenesis, trafficking) and tumor intrinsic pathway genes driving the TiME phenotyping (Fig 2E) were also investigated.

**Figure 2.**
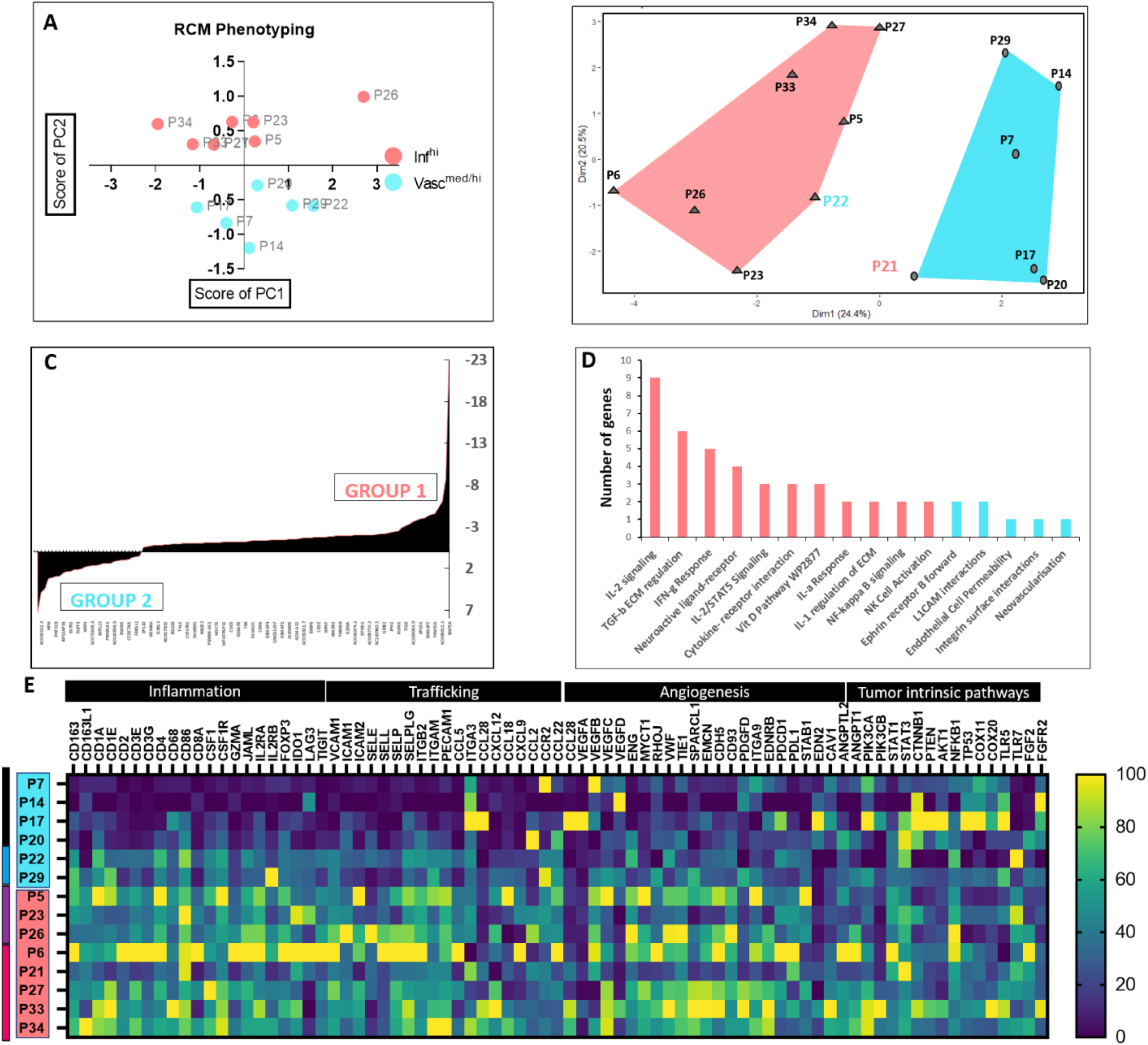
Higher immune signaling and activation in Inf^hi^ phenotypes as compared to Vasc^hi^ phenotypes. **A.** *In vivo* TiME phenotypes in 14 patients grouped in Inf^hi^ and Vasc^med/hi^ for gene expression analysis. **B.** k-means clustering on CIBERSORT output to characterize immune cell abundance reveals two groups similar to RCM phenotypes with two misclassifications (P21 and P22). **C.** DGEA on ~30,000 genes was performed for Inf^hi^ and Vasc^med/hi^ TiME phenotypes:114 differentially expressed genes were found, of which 85 genes were upregulated in Inf^hi^ and 29 genes in Vasc^med/hi^. **D.** Relevant gene ontology and pathway analysis on the differentially expressed genes (p<0.05). **E**. Heatmap of differences in major inflammation, angiogenesis, trafficking and tumor-intrinsic pathway gene expression is shown. The 4-group phenotype is represented by the color bar on the left. RCM: reflectance confocal microscopy, Inf: inflammation, Vasc: vascular; DGEA: differential gene expression analysis

Next, we linked patient-specific responses to topical immunotherapy with TiME features and phenotypes. Within this pilot cohort, most responders (4 out of 7) belonged to the Vasc^lo^Inf^hi^ phenotype (Fig 3A). Differences in TiME features between responders and non-responders suggest high vessels and stromal macrophages/dendritic cells in non-responders (Fig 3B), also seen in representative RCM images (Fig 3C-D). Higher leukocyte trafficking, stromal vessels, and stromal macrophages were present in 50%, 100% and 86% of the non-responders. Additional analysis for modeling TiME features and tumor phenotypes with response to treatment was performed. Linear separability plots confirmed stromal vessels and stromal macrophages variables led to high separation between responders and non-responders (Fig S6A). Linear regression models using specificity or Akaike Information Criterion (AIC) coefficient as outcome measures (Fig S6B) demonstrate low predictive power of inflammation as a variable, either as “tumor-infiltrating lymphocytes” or “intratumoral inflammation” with accuracy of 46% and 61%, respectively (Fig S6C). Addition of stromal vessels to intratumoral inflammation or tumor-infiltrating lymphocytes (Fig S6C) as features in the linear regression model resulted in best model performance (71% sensitivity, 83% specificity and 76% accuracy).

**Figure 3.**
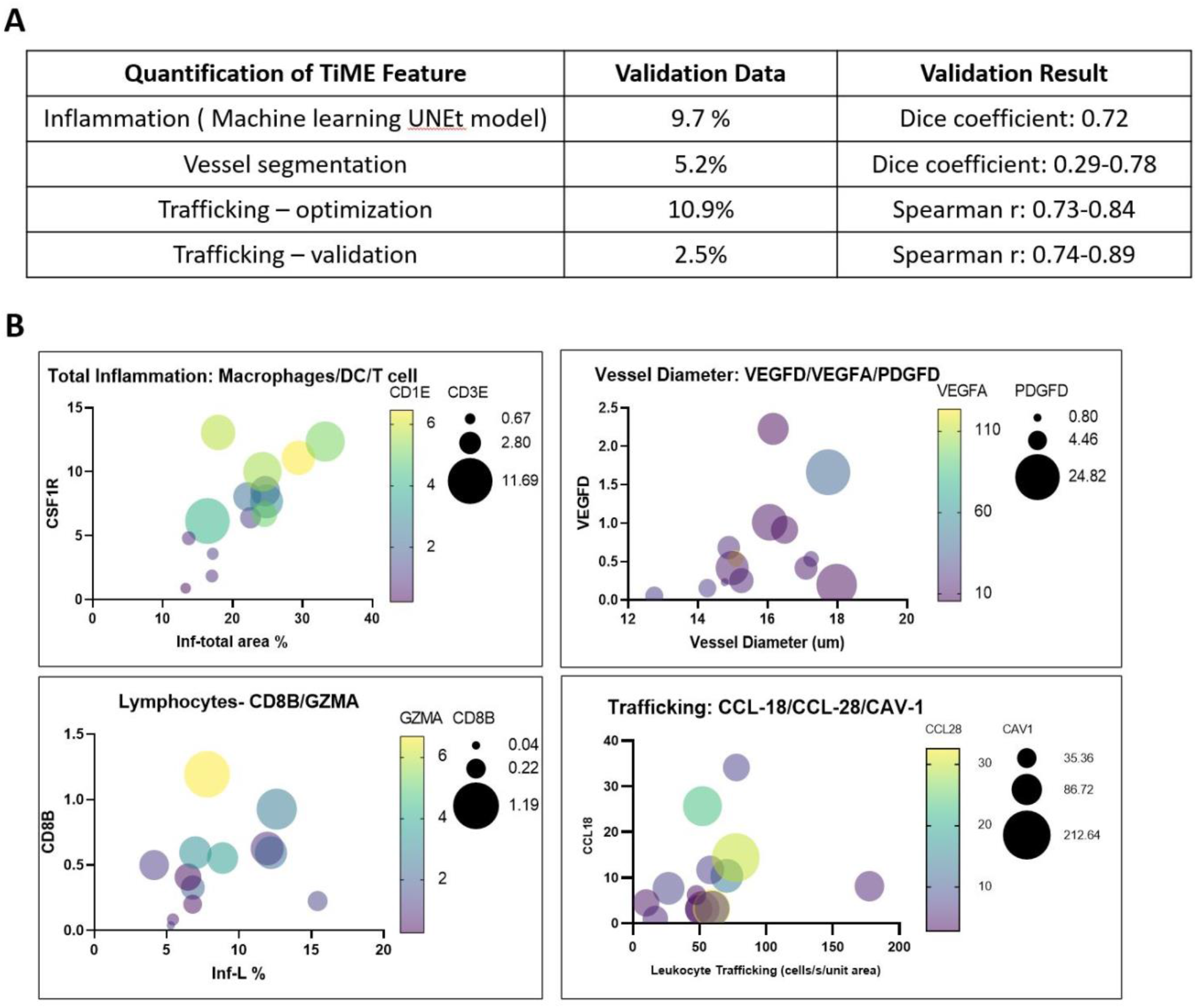
Quantification of TiME features and correlation with gene expression. **A.** Summary of results for validation performed after each automated quantification. **B.** Spearman correlation for TiME features and corresponding gene expression: total area % inflammation with CSF1R (r=0.73, CI: 0.32 to 0.91), CD1E (r=0.64, CI: 0.15 to 0.87) and CD3E (r=0.51, CI:−0.04 to 0.82), between total round-ellipsoid leukocytes area % with CD8B (r=0.6, CI: 0.1 to 0.86) and GZMA (r=0.53, CI: −0.01 to 0.83), vessel diameter with VEGF-D (r=0.459, CI: −0.1 to 0.80), VEGF-A (r=−0.471, CI: −0.81 to 0.09) and PDGFD (r=0.538, CI: 0 to 0.84), and trafficking with CCL-18 (r=0.561, CI: 0.02 to 0.84), CAV-1 (r=0.46, CI: −0.1 to 0.86) and CCL28 (r=−0.42, CI: −.016 to 0.78). p values were significant for all comparisons except CCL28. CSF1R: colony stimulating factor 1-receptor; CD: cluster of differentiation; GZMA: granzyme A; VEGF: vascular endothelial growth factor; PDGFD: platelet derived growth factor D; CCL: CC-chemokine ligand; CAV: caveolin.

Finally, we investigated automated quantification of RCM TiME features— immune cells, leukocyte trafficking and vascular features using machine learning and image processing algorithms to enable objective quantitative comparisons between RCM TiME features and biological markers. Quantification of immune cell density was explored using a machine learning segmentation model, U-Net. Representative images and segmentations are shown in Fig S7A. Image-processing algorithms were used for quantification of vascular features (vessel area, diameter and number) and leukocyte trafficking (Fig S7B-G). Each analysis was validated on a subset of data (Fig 4A). Correlation between manual evaluation and automated quantification was also evaluated (Fig S7H). Subsequently, correlation of RCM TiME features with corresponding gene expression for inflammation, angiogenesis and trafficking were also computed (Fig 4B).

**Figure 4.**
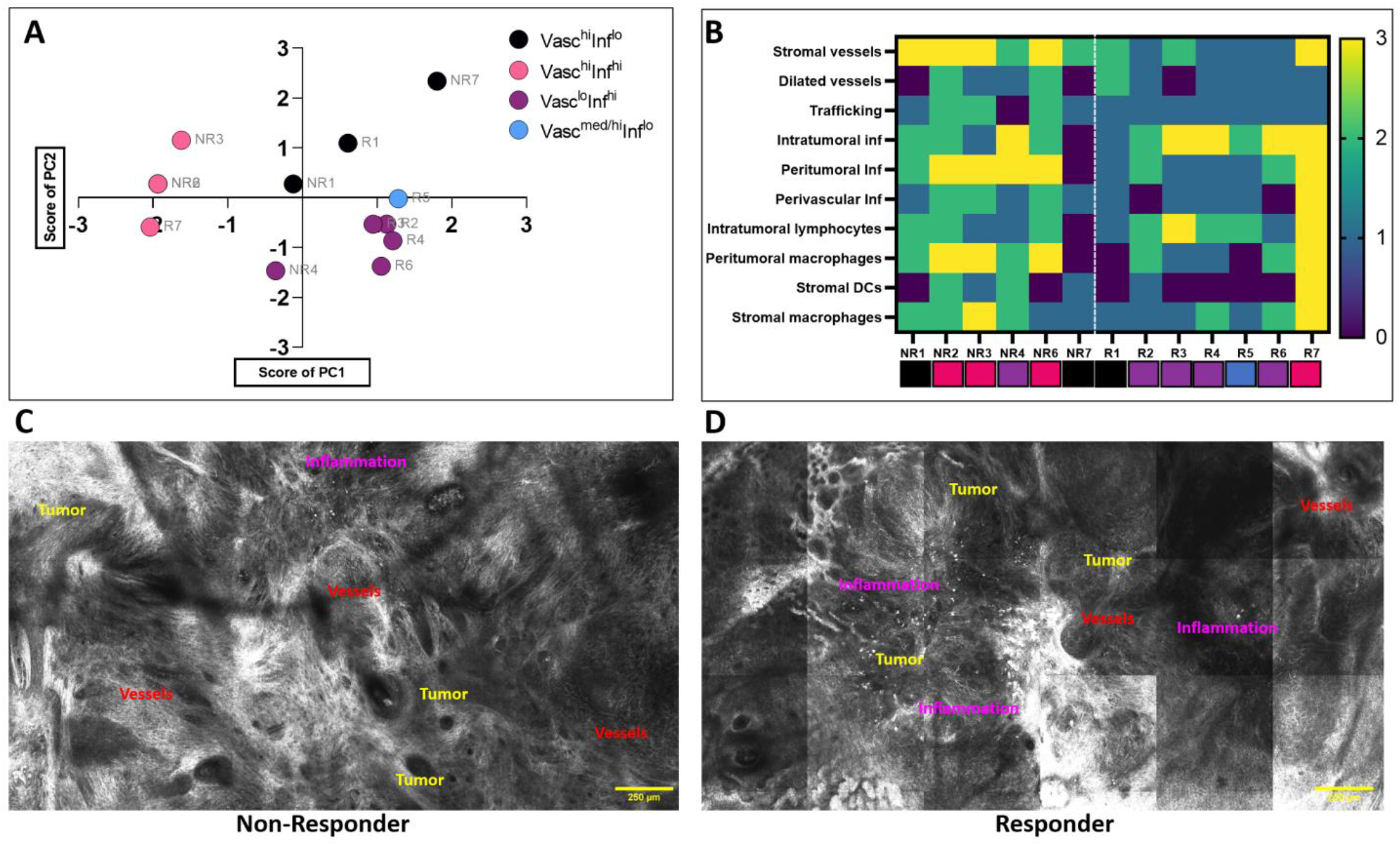
TiME phenotype and RCM features correlate with treatment outcomes. **A.** PCA on TiME features of patients receiving imiquimod and confirmed as responder or non-responder based on tumor clearance on RCM imaging. **B.** Heatmap of differences in RCM TiME features of responders and non-responders. **C, D**. Representative TiME features in responders and non-responders demonstrate greater number of vessels in the stroma, immune excluded tumors and inflammation in the stroma in the non-responder image while higher intratumoral and stromal inflammation (dendritic cells, macrophages, lymphocytes) with comparatively less number of vessels in the responder image. PCA: principal component analysis; TiME: tumor-immune microenvironment; RCM: reflectance confocal microscopy.

## Discussion and Conclusion

Our findings indicate that *in vivo* TiME phenotyping show correlation with underlying immunological and vascular states and treatment responses to topical immunotherapy. We show that TiME features on RCM can be reproducibly evaluated and correlated with histopathological TiME features. The TiME phenotypes correlated with presence of CD3^+^ T cells and TLS area, and Inf^hi^ was characterized by more inflammation attributes as compared to Vasc^med/hi^. Of the thirteen patients who underwent prospective treatment with topical immunotherapy, 7 patients responded to the immunotherapy treatment while 6 were non-responders. On this small cohort of patients, we identified the importance of additional TiME features in mediating response to immunotherapy and correlated Vasc^lo^Inf^hi^ phenotype with maximal response. Investigations into specific genes and pathways in individual patients demonstrate high immune and angiogenic signatures, including features characteristic of inflamed phenotypes such as CXCL9 (T-cell recruitment inside tumors)^34^, induction of suppressor pathways PD-L, FOXP3, IDO-1 and exhaustion markers TIGIT^35^ and higher memory cells (associated with improved overall survival^36^) in Vasc^hi^Inf^hi^. This phenotype likely had normalized vasculature, as seen in high expression of VCAM1, ICAM1, L-selectin, CCL2 and higher intratumoral inflammation^18,20^. Conversely, Vasc^hi^Inf^lo^ show features characteristic of endothelial anergy and immunosuppression (poor immune infiltration in tumors), including downregulation of major adhesion molecules (VCAM, ICAM1, ICAM2, SELL), higher VEGFD, relatively higher EDN2 and ITGA3. While VEGFD is implicated in blood and lymph vessel dilation^37^, genes such as CCL-22, CCL-28, ITGA3, may likely be compensating for decreased VCAM1, ICAM1in patients with higher trafficking^38^. Higher CCL28 correlation was seen only in the Vasc^hi^ phenotype with high trafficking (data not shown), suggesting the possible role of CCL-28 in trafficking of T-regulatory cells in these patients^39,40^. This effect could be potentiated by upregulation of immunosuppressive tumor-intrinsic factors (CTNNB1, PTEN, COX11)^41,42^ likely inducing immune exclusion and endothelial anergy. Alternatively, Vasc^lo^Inf^hi^ corresponded to a highly inflamed phenotype with lower immunosuppressive vascular features with higher intratumoral inflammation in RCM. This feature was also reflected in the prospective imiquimod study where highest proportion of responders were found to belong to the Vasc^lo^Inf^hi^ phenotype. Relatively higher expression of TLR7 (agonist of Imiquimod) in this phenotype (Fig 2E) could also indicate higher Imiquimod response in this phenotype. Additionally, immunophenotyping to correlate RCM phenotypes in a pilot study on 3 BCC tumors indicated higher activated CD8^+^GZMb^+^ and CD8^+^Ki-67^+^ cells in a patient with features of Inf^hi^ as compared to a patient with Vasc^hi^ also suggesting the inflamed nature of Inf^hi^ phenotype (Fig S8).

Thus, these results suggest the feasibility of identifying not only the TiME phenotypes, but also the mechanism of immunosuppression and lack of response that can be exploited for treatment of cold, non-inflamed and non-responsive tumors. Anti-angiogenic agents can overcome endothelial cell anergy and reinduce adhesion molecule expression, resulting in increased leukocyte infiltration into tumors^18^. Thus, skin cancer phenotypes with the Vasc^hi^Inf^lo^ phenotype could benefit from additional synchronous treatment for vessel normalization through pharmacological targeting of the WNT/β-catenin pathway or anti-angiogenic topical treatments (COX-2, basic fibroblast growth factor or bFGF inhibitors) towards treatment optimization^40,43^. Additionally, immune-cell infiltration into tumors does not necessarily warrant response, thus converting excluded anergized tumors into inflamed may fail as a treatment strategy^15^. Longitudinal and non-invasive monitoring of the treatment-induced alterations can help assess response and uncover resistance mechanisms, elucidate newer TiME targeting therapies and achieve further treatment optimization.

Predicting response/resistance immunotherapy as well as understanding how tumors escape host immune defenses can facilitate effective treatment strategies. Currently known predictive mechanisms including the IFN-G gene panel and the “Tumor Inflammation Signature” have shown some promise in predicting response to immune checkpoint blockade^44,45^. A more comprehensive and quantitative analysis of major determinants of anti-tumor immunity within the TiME will improve patient stratification, which will ultimately improve current cancer management paradigms and introduce a more personalized immunotherapy platform, similar to the IMPACT^TM46^. *In vivo* imaging is crucial for studying these dynamic interactions within TiME, since active vascular processes such as leukocyte trafficking are optimally studied as a live dynamic process, as opposed to *ex vivo* tissue studies on vasculature which show inconsistent vessel measurements^23^. Using RCM and skin cancers as a model, we demonstrate the proof-of-concept for unperturbed characterization and TiME phenotyping inside patients. Furthermore, feasibility of automated quantification to generate stronger quantitative correlations was also demonstrated.

While this study demonstrates a combination of high-resolution spatially resolved and dynamic imaging, limitations included grayscale-limited specificity tissue contrast and imaging depth to 0.2-0.25 mm. The label-free approach enables visualization of all TiME features, but is limited in specificity for functional phenotyping. This potential limitation may be overcome by monitoring longitudinal spatio-temporal changes in immune cells repeatedly over prolonged periods. The limited depth of imaging fails to capture deeper TiME features. With the current state of RCM devices and technology, this approach is currently restricted to accessible diseases and cancers, namely skin cancer, head-neck cancer, cervical cancers, cutaneous lymphomas, cutaneous metastasis. In the future, extensive validation with targeted molecular correlations on precision biopsies^47^ will enable better correlations. Exhaustive molecular validation using flow cytometry and single-cell RNA-seq and spatial transcriptomics^48,49^ on subsequent models will facilitate improved understanding of RCM phenotyping. Complementary multimodal approaches^50^ such as dynamic optical coherence tomography or optical frequency domain imaging for imaging vasculature, lymphatics and tissue viability ^51,52^, multiphoton microscopy for better contrast and collagen delineation^53–55^, photoacoustic microscopy for functional vascular imaging^56^ and fluorescence lifetime imaging for probing immune cell specificity and activation states^57^ may be necessary to further enhance *in vivo* TiME visualization and enhance current TiME phenotyping. Through robust prospective studies, fundamental basis of phenotyping and their correlation with variable treatment responses in cancer immunotherapy systems will be explored for better patient stratification. These initial findings will enable hypothesis-driven research for developing novel druggable targets and gaining mechanistic insights on host anti-tumor immune response in various bedside cancer settings in human patients.

## Methods

### Patient recruitment and imaging

Patients referred for physician consultation, Mohs surgery or wide local excision at Memorial Sloan Kettering Cancer Center (MSKCC), NY were prospectively enrolled for this study under MSKCC-IRB approved protocols. Patients (aged 18 or older) with either a previously biopsied or clinically suspected keratinocytic [basal cell carcinoma (BCC), squamous cell carcinoma (SCC), actinic keratosis(AK)], melanocytic (melanoma) lesion or drug rash that were amenable for imaging were accrued consecutively at Memorial Sloan Kettering Cancer Center (MSKCC) after written informed consent. Patients (n=9) with suspected BCC (n=13) and selected for topical immunotherapy Imiquimod were also enrolled.

### In vivo imaging

*In vivo* RCM imaging was performed prospectively on 105 lesions (or cases) using an RCM (either VivaScope 1500 or a handheld VivaScope 3000, Caliber I.D., Rochester, NY) and/or an integrated handheld RCM-OCT prototype. Images were acquired and interpreted in real-time at the bedside to select representative areas with tumor, immune cells and blood vessels across the lesion by 2 investigators (M.C. and A.S.) having more than 4 years of RCM reading experience. Mosaics (large area sampling), stacks (depth sampling), scanning and single field-of-view (FOV) videos were acquired from multiple regions within the lesion and saved in an online database (Vivanet, Caliber ID, Rochester, NY) or on a local drive. Individual images (0.75 × 0.75 mm) from stacks and temporal single FOV frames were used for automated quantification of immune cells, and vascular features, respectively.

### Patient tissue

Biopsies (targeted or non-targeted) taken as standard-of-care or for research use were used for histopathological, immunohistochemical, RNA-sequencing and flow cytometry correlations. Formalin-fixed paraffin embedded (FFPE) specimens from 34 lesions (33 for main study, 1 for cell identification in Fig S1) were used for histopathological and immunohistochemical correlations. Eight tissue sections were provided by a collaborator for tertiary lymphoid structure studies. RNA- extraction was on 25 lesions and RNA-seq was performed on 14 lesions. Imaging-guided targeted biopsy was performed on 6 lesions: frozen sections followed by pathology/IHC on 3 lesions and flow cytometry was performed on 3 lesions.

### RCM Data

Stacks and videos from 97 lesions were analyzed in this study. Machine learning-based immune cell quantification was explored on 1026 frames from 92 cancer lesions. Each case contributed 5-27 independent images. The algorithm was tested on 652 independent images from 33 cases at ~20 representative images/lesion. For vascular feature quantification, 438 single FOV videos (39, 813 frames) from 48 cases were selected. Each lesion contributed 1-31 videos. Quantified values from 270 videos in 26 BCCs in the analyzed set were used for subsequent correlations with manual evaluation and gene expression.

### RCM manual evaluation

RCM features were manually evaluated (Fig S1A-D) by 4 readers with at least 4 years’ experience (AS) or >20 year experience in interpreting RCM images for the main study on 33 cases (MC, SG) or treatment response study on 13 cases (CMAF). The major features evaluated on manual reading included number of vessels, dilated vessels, trafficking, intratumoral inflammation, peritumor inflammation and perivascular inflammation. These features were graded on a scale of 0-3 after exhaustive review of data from each patient. For imiquimod response study, spatial distribution of vessels and type of three immune morphologies (dendritic cells, leukocytes and macrophages), mucin and tumor regressing areas were noted for more comprehensive assessment and correlation with treatment response.

### Histopathological evaluation

Same TiME features evaluated on RCM were also graded on digitized histopathological slides of 33 patients by a board-certified dermatopathologist (MG).

### Agreement and correlation studies

Agreements between two readers’ manual evaluations for binary RCM feature presence were quantified using Cohen’s kappa coefficients. For the evaluations between RCM and histology, agreement regarding the extent of each feature presence was quantified using linearly weighted Gwet’s AC1 for each of the two RCM readers to the single histology reader. The simple average of the two Gwet’s AC1 scores were reported for each feature. Binary feature presence on RCM versus histology was derived the same way after recoding the manual evaluations.

Correlations between automated quantified and manual features were computed using Spearman’s correlation (confidence interval: 95%). Spearman correlation between RCM quantified features and immune-related, trafficking-related and angiogenesis genes were also computed. .

### Statistical clustering for TiME phenotyping

Unsupervised statistical clustering on TiME features was performed to explore classification trends or phenotypes. Principal component analysis (PCA) was used for analysis of 6 manually evaluated RCM TiME features—intratumoral, peritumoral and perivascular inflammation, vessel area/number, dilated vessels and trafficking. Clustering was performed on principal components values. Centered method (data scaled such that mean =0, SD unchanged) was used for the PCA. The number of PCs were selected such that largest eigenvalues together accounted for 95% of the total variance. Loadings are presented along with scatter plot.. For PCA on 27 patients, the largest absolute loadings (PC1: dilated vessels, trafficking and PC2: peritumoral, perivascular inflammation) were used to establish the phenotypes. Patients with negative or small values of PC1 and PC2 were denoted as Vasc^hi^Inf^hi^, patients proximal to the PC1 loadings were denoted Vasc^hi^Inf^lo^ while distal to PC1 loadings were Vasc^med/hi^Inf^lo^, and patients distal to main PC2 loadings but proximal to intratumoral (IT) inflammation loading vector were denoted as Vasc^lo^Inf^hi^. The ‘hi’ and ‘lo’ indicate the prevalence of the feature in the phenotype, high corresponds to manual score of 2 or 3, low to 0 or 1 (on a scale of 0-3, Fig S1D) while ‘med’ denotes medium feature prevalence (between 1-2). The Vasc^hi^Inf^hi^ phenotype was characterized by higher peritumor and perivascular inflammation while the Vasc^lo^Inf^hi^ was characterized by higher intratumoral inflammation and lower peritumor/perivascular inflammation. PCA for individual gene groups (inflammation^30^, angiogenesis^28^ and trafficking(GO Pathways^58^)) were also analyzed to derive correlations with RCM phenotypes (Fig S2E). Another PCA on response to imiquimod was analyzed using same parameters (Fig 4A). PCA was performed in Graphpad 9.0.

### Immunohistochemistry

CD3^+^, CD1α, CD68^+^ and CD20^+^ IHC were performed on BOND RX(Leica). The protocol for the Bond Rx platform included ER2 (High pH buffer) −30 minutes for Heat retrieval followed by 30 minutes incubation time for Primary Abs (Santa Cruz Biotech, US). Polymer Detection was through DAB Kit (catalog DS9800). For the dual CD3/CD20 sequential stain, ER2 −30 minutes for heat retrieval, 30 minute incubation time for Primary Ab followed by Polymer Detection kit. This was followed by ER2-20 minute, 30 minute -incubation with second Ab, Polymer refine Red detection KIT, (catalog # DS9390).

### IHC evaluation and quantification

CD3^+^, CD20^+^ stains were evaluated on 33 cases by a board-certified dermatopathologist (MG) for presence/absence of ulceration/erosion and CD3^+^ T-cells, CD20^+^ B-cells, total lymphocytes (CD3^+^ CD20^+^). In addition, for each immune marker, features were evaluated on a scale of 0-3 where 0 is absent and 3 is highest. These features included predominant distribution, TILs, trafficking and distribution at tumor periphery. Tertiary lymphoid structures (TLS) labeled by dual CD3^+^/CD20^+^ staining on 40 cases were also analyzed for total TLS numbers, TLS dimensions (maximum dimensions in × and Y) and tissue size (maximum dimensions in × and Y) to compute TLS numbers/mm2 and TLS area coverage/mm^2^. Within defined TLS and non-TLS areas (used as control), tumor killing as defined on histopathology was noted, and TILs in TLS-adjacent tumor nests were counted. For TLS positive patients, both TLS and non-TLS areas were evaluated for TILs and tumor killing, in TLS negative patients, TILs and tumor killing were specified in defined areas (Fig S3). Two-sided confidence intervals for median proportion of TLS area coverage was derived from percentile bootstrapping approach. Mann-Whitney U tests (two-sided) were used to quantify the statistical significance in differences between median proportion of TLS coverage across binary clinical factors such as ulceration presence and NMSC classification. Generalized estimating equations (GEE) were used to estimate association between local TiME TLS presence with both TILs presence and tumor killing by clustering on histologic specimen using an exchangeable correlation structure. This approach was applied to the binary classification of local TLS area versus local control area to the continuous outcome of local TILs cell count density. GEE was applied exclusively to TLS regions to model the local TLS area coverage to both the continuous outcome of TILs cell count using Gaussian link function and to the binary outcome of tumor killing using the logit link function.

### RNA analysis on GEO datasets

Two previously-published RNA-seq data sets (GSE125285, GSE128795) and one microarray data set (GSE53462) for Basal Cell Carcinoma samples were downloaded from the Gene Expression Omnibus (GEO). GSE125285 and GSE128795 contained pre-processed RNA-seq data, however, the microarray data was indexed to Illumina Probes^59,60^. First, those probes with high detection p-values (p > 0.05 for 13 out of the 26 samples) were filtered out, leaving 23, 176 probes remaining from an initial value of 47, 323. ProbeIDs were matched to common gene identifiers using illuminaID2nuID^12^. Of the remaining probes, 5, 778 did not have a unique gene associated with them. We took the value with the highest expression to have each gene represented only once, leaving 17398 genes. We created phenotype groupings a priori via unbiased clustering through immune-related genes provided by Nanostring^61^©. For the whole transcriptome analysis, we used the built-in R heatmap function (stats 4.0.2) to create phenotype clusters. The heatmap revealed 2 groups on which the DGEA analysis was performed. Functional enrichment analysis was performed using GO enrichment analysis (https://go.princeton.edu/tmp/5497206//query_results.html), and each enriched ontology hierarchy (false discovery rate (FDR) < 0.05) was reported with two terms in the hierarchy: (1) the term with the highest significance value and (2) the term with the highest specificity.

### RNA extraction

FFPE sections were deparaffinized using the mineral oil method. Briefly, 800μL mineral oil was mixed with the sections and the sample was incubated at 65°C for 10 minutes. Phases were separated by centrifugation in 360μL Buffer PKD and Proteinase K was added for digestion. After a three-step incubation (65°C for 45’, 80°C for 15’, 65°C for 30’) and additional centrifugation, the aqueous phase containing RNA was removed and DNase treated. The RNA was then extracted using the RNeasy FFPE Kit (QIAGEN catalog # 73504) on the QIAcube Connect (Qiagen) according to the manufacturer’s protocol with 285μL input. Samples were eluted in 13μL RNase-free water.

### Transcriptome sequencing

After RiboGreen quantification and quality control by Agilent BioAnalyzer, 356-500ng of total RNA with DV200% varying from 88-93 underwent ribosomal depletion and library preparation using the TruSeq Stranded Total RNA LT Kit (Illumina catalog # RS-122-1202) according to instructions provided by the manufacturer with 8 cycles of PCR. Samples were barcoded and run on a HiSeq 4000 in a PE100 run, using the HiSeq 3000/4000 SBS Kit (Illumina). On average, 78 million paired reads were generated per sample and 20% of the data mapped to mRNA.

### CIBERSORT analysis

CIBERSORT was used for the immune cell analysis to delineate immune subsets using 584 genes for 22 immune cell types^29^. Transcript per million values were used as input. CIBERSORT chooses the record with the highest mean expression across the mixtures during analysis. The gene expression file with 14 cases was uploaded to CIBERSORT as a mixture file, and CIBERSORT was run with the following options: relative and absolute modes together, LM22 signature gene file, 100 permutations, and quantile normalization disabled. Sample distance matrix resulting from immune cell distribution, k-means clustering and differential gene expression analysis (DGEA) were used to interpret CIBERSORT output.

### Differential gene expression analysis (DGEA)

DESeq2 (ver 1.28.1) was used to perform differential gene expression analysis comparing RCM groups 1 vs 2. Genes with an absolute log2 fold change of >= 0.5 an adjusted p-value of < =0.1 were considered significantly changed. Log transformation was then performed on the full gene expression matrix with the *rlog* function and the transformed read counts of the 114 significantly changed genes were extracted for unsupervised hierarchical clustering analysis with pheatmap (ver 1.0.12, clustering_method = “complete”)

### Gene set enrichment analysis

471 angiogenesis genes were identified to generate the angiogenesis core gene set and 547 immune genes were extracted from CIBERSORT analysis as mentioned above. Top 10% genes differentially expressed between the RCM groups 1 vs 2 ranked by absolute fold change were identified and ranked from the highest to the lowest fold change. Gene set enrichment analysis was then performed on the 2,529 genes to calculate the enrichment score for the angiogenesis and immune gene sets with the R package fgsea (ver 1.14.0) and the *fgseaMultilevel* function.

### Response to Immunotherapy analysis

Correlation of TiME features and phenotypes with response to topical immunotherapy Imiquimod were analyzed on 13 cases. The patients were imaged at baseline (T0) and the TiME features were analyzed with respect to response to treatment. Linear regression modeling was undertaken to quantitatively identify the predictor variables for response to imiquimod and compared against the known “standard” which is tumor-infiltrating lymphocytes and intratumoral inflammation. In order to measure the predictive power of each feature, we trained predictive models in a leave-one-out cross-validation fashion and measured the model performance by inferring on the left-out test sample (out-of-bag estimates). This procedure was followed in an iterative manner, where we selected a single feature that gives the highest performance and added a new feature that provided the highest performance in each iteration. Model performance was measured by calculating specificity (higher the better) on the out-of-bag estimates and Akaike Information Criterion (lower the better) value of the model. In this way, the features were prioritized according to their predictive power. The best performance among the AIC prioritizing models was 85% sensitivity, 66% specificity and AIC=−15.06 with 8 variables while the best performance among the specificity prioritizing model was 71% sensitivity, 83% specificity and AIC= −15.06 with 13 variables. Three of eight features (stromal vessels, peritumoral vessels and stromal macrophages) overlapped between the two models. Moreover, we also examined the linear separability of (i) individual features by looking at the histogram of feature values for each sample, and (ii) each pairwise feature combination by examining kernel density estimation plots. PCA on these patients was also performed, as described above.

### Immunophenotyping

Freshly excised 3 mm punch biopsies from 3 BCC lesions were stored in DMEM media for 24-48 hours at 4 °C. Tissue were transported to cell culture lab on ice. Cell suspensions were generated according to the following protocol^62^. Cells were processed for surface labeling with anti-CD3, anti-CD45, anti-CD4, and anti-CD8 antibodies. Live cells are distinguished from dead cells by using the fixable dye eFluor506 (eBioscience). They were further permeabilized using a FoxP3 fixation and permeabilization kit (eBioscience) and stained for Ki-67, FoxP3, and Granzyme B. Data were acquired using the LSR II flow cytometer (BD Biosciences). Data were analyzed with FlowJo software (Tree Star)^63^.

### Quantification

#### Machine learning for immune cell

A pixel-wise segmentation model was trained for 4 different morphological patterns (dendritic cells, macrophages, leukocytic round-ellipsoid cells and miscellaneous immune cells) imperative for TiME analysis. We binned them into 2 classes as class 1: Dendritic cells and Melanophages (macrophages), Class 2: round-ellipsoid leukocytes and miscellaneous immune cells. As a third class we also labelled areas that did not contain any of these patterns as background. 1026 RCM images from 93 lesions were labelled pixelwise for these 3 classes in a non-exhaustive manner, where we only labelled examples of these patterns (Fig S7A). A total of 9% of the pixels were labelled (6% Class 1, 3% Class 2 and 91% Class 3). We trained a 3 class UNet^64^segmentation model using the MONAI framework^65^. We used 926 images for training and 100 independent images for testing the model. Based on our former studies^66,67^, we downsampled the RCM images to 256 by 256 pixels (corresponding to 2 μm resolution) for the sake of computational efficiency. We use a learning rate of 5e-2, batch size of 64, and SGD optimizer with Nesterov momentum. We also used image augmentation such as random rotation, flipping, elastic-affine deformation, intensity scaling, to increase the training dataset size. The model is trained for 90 epoch using DICE loss. After 90 epochs we did not see any improvement in the loss. Dice similarity coefficient of 0.72 was found for these 3 classes (Fig 4A).

#### Vascular features

For all video frames, a two-step image stabilization procedure was used to remove the significant motion found in each movie segment. Firstly, a linear pre-alignment is performed to minimize large scale motion in Fiji^68^ using the SIFT feature plugin Plugins->Registration-> Linear Stack Alignment with SIFT and default parameters. Stabilized images are then automatically cropped in Matlab (mathworks.com) to remove black background and include only areas within the field of view during the entirety of the movie segment. The crop rectangle is computed automatically by iteratively removing the row or column of pixels which contains the most blank pixels in a temporal min image until all outer edge rows and columns that contain more than three quarters blank pixels are removed. A second custom nonlinear stabilization was then performed in Matlab to remove large scale tissue deformations over time. Frame t+1 first has its histogram equalized to match frame t and then is aligned to frame t using the imregdemon procedure with four pyramid levels and steeply decreasing iterations of alignment at successively finer scales (iterations, [100,50,10,1]). Frame t+2 is then aligned with the transformed frame t+1 and so on. Cropping of all regions not in view throughout the movie is again performed via the same procedure.

#### -Trafficking

##### Background Subtraction

A background image is estimated for each frame as the median per pixel over a temporal window of 6 s centered on the current frame. Where movie temporal resolution differs, the window in frames is adjusted accordingly. This background estimate is subtracted out of the current frame, largely isolating moving cells on a dark background. We experimented with sparse linear methods for background subtraction, but found increased distortion in extracted foreground cells were a persistent problem across methods (data not shown). Mean and min background estimates were also tried, as well as dividing through by, rather than subtracting, background estimates, which desirably enhances dim cells. This advantage was offset by noise enhancement in non-vessel voids in the tissue (data not shown).

##### Tracking

Background subtracted images are exported from Matlab as 32bit OME tiffs and imported into Fiji. Tracking is then performed in Trackmate^69^ using DoG spot detection (subpixel=true; radius=7.5pixels (7.5/1.33= 5.63 micron); threshold=1.6) and the LAP tracker with no splitting, merging or gap closing, and a max match distance of 20 pixels (20/1.33 = 15.03 micron). The tracklets found are then filtered in Matlab to remove spurious tracklets corresponding to imperfectly removed background elements (this occurs particularly during changes in z during imaging) or tracks strung together from different fast moving circulating blood cells while preserving the desired target population. Features used to measure tracklet desirability are detailed below. Thresholds were set quantitatively and automatically to maximize correspondence between automated results and manual counts on an initial training set of 40 movies (approx. 10% of overall data). Three different temporal windows ranging from 0.6s, 0.8 s and 1s were investigated for total quantification of rolling, crawling and adherent cells. Constrained optimization within a restricted range was adopted, although fully independent threshold optimization was also investigated (Fig S7C-F). Moderate-high correlation (0.79-0.82) was observed during first optimization following which trafficking was quantified on remaining videos. Final validation using manual counts on a subset of videos (~2.5% of total data) by two readers with high inter-reader concordance found high correlation (0.74-.0.89) for different temporal windows (Fig S7G). Temporal window 3 was selected for subsequent analysis to ensure inclusion of especially faster trafficking processes (rolling cells) in shorter blood vessels. The correlation was worse for videos with remnant motion after two-step motion minimization strategy, suggesting need for minimizing in axial and lateral motion during data acquisition, and use of more efficient motion removal algorithms in future.

The Tracklet Parameters used are as follows:

Displacement=[15.41,16.92,16.92]um([20.5,22.5,22.5]px)

Consistency=[58,58,58] degrees

Quality=[1.6,1.65,1.75] arbitrary units;

Length=[0.6 s, 0.8 s or 1 s])

Where,

Displacement- total displacement between tracklet start and end point, in pixels (tracklets with lower displacement are discarded)

Motion Consistency – average angle between the motion vector of the track at successive timepoints in degrees (tracklets with higher angular difference are discarded)

Quality- average quality of detections making up the tracklet as measured by Trackmate (lower average quality tracklets are discarded)

Length – duration in s of tracklet, in all cases this was set to the thresholds used in manual counts (shorter tracklets are discarded)

#### -Blood vessel segmentation

Manual segmentation of blood vessels was performed using an open-source segmentation platform called 3D Splicer (https://www.slicer.org/)^70^ on 25 randomly selected videos. Two videos were discarded from analysis due to extreme Z-motion. The remaining 23 videos were processed to display only every 10^th^ frame to mimic the automated segmentation approach; each frame in the resulting file was segmented. The entire video segmentation was exported as a Nifti (.nii) file format and imported into Matlab as a 3D image array, where consecutive images in the array correspond to consecutive frames in the RCM video. Ensuring that the consecutive frames are registered, our assumption for detecting the vessels was that the areas of high variation between consecutive frames correspond to vessels. In order to suppress the variation due to speckle noise in the RCM images, we first applied a gaussian smoothing filter (sigma = 1px). Then we applied a finite impulse response high pass filter (F = [0.5,−1,0.5]) and smooth out the extracted pixel-wise variation in time using a 7-by-7 median filter. We then subtract the mean variation of each frame to eliminate the slowly varying areas, and obtain a variation map for the whole video by accumulating the variation over the entire video sequence. We finally apply otsu thresholding the final variation map to find the areas of vessels in the videos. To smooth the border of the vessels and clean out the noise in the segmentation, we applied morphological closing operation on the binary segmentation map and clean segmented areas smaller than 0.1% and larger 10% of the entire frame. Dice similarity coefficients were calculated for comparing manual and automated vessel segmentation (Fig S7B, 4A).

## Acknowledgement

We would like to acknowledge Dr. Anjali Rajadhyaksha for assistance with experimental planning and scientific discussion. We also acknowledge MSKCC Cores: Integrated Genomics Operation Core, funded by the NCI Cancer Center Support Grant (CCSG, P30 CA08748), Cycle for Survival, and the Marie-Josée and Henry R. Kravis Center for Molecular Oncology, Molecular Cytology Core Facility, Pathology Core, Flow Cytometry Core. In addition, Ms. Cassidy Cobbs, Ms. Marina Asher, Mr. Afsar Barlas and Mr. Eric Chan for assistance with experimental planning, data analysis and immunohistochemical staining. We also acknowledge our funding sources NIH/NCI Cancer Center Support Grant P30 CA008748 (MSKCC) and NCI/NIBIB R01EB020029 (Milind Rajadhyaksha), Melanoma Research Alliance (Aditi Sahu) and the Chan-Zuckerberg Initiative (Anthony Santella).

## Data availability statement

The datasets generated during the current study are available from the corresponding author on reasonable request. The RNA-expression datasets generated and/or analyzed during this study will be made available on gene expression omnibus (GEO).

## Code availability statement

Custom image and data analysis scripts for Fiji and Matlab developed for quantification of imaging data are available on https://github.com/mskccmccf/TiME-analysis

