## Supplementary Table and Figures for "Cellular-level phenotyping of tumor-immune microenvironment (TiME) in patients in vivo reveals distinct inflammation and endothelial anergy signatures"

**Supplementary Table/Figures:**

| Cell | Cell Size (μ) | Shape | Nuclear Size (μ) | Nuclear Shape | Refractive Index | Scattering Coefficient | References |
|---|---|---|---|---|---|---|---|
| **T Cell** | 8-10 | Round | 6-7 | Irregular circular shape | 1.36 (1.34-1.37) | $0.54*10^4$ (STD=$0.076*10^4$) | [1], [2] |
| **B-Cell** | 8-10 | Round | 6-8 | Nucleus indented | 1.36 (1.34-1.37) | $0.58*10^4$ (STD=$0.075*10^4$) | [3], [2] |
| **Neutrophil** | 12-14 | Irregular-round | 3-4 | Multilobed nucleus | 1.4 (1.35-1.42) | $4*10^4$ (STD=$0.57*10^4$) | [3,4], [2] |
| **Eosinophil** | 12-15 | Round | 3-4 | Bilobed | 1.4 (1.35-1.42) | $4.6*10^4$ (STD=$1.04*10^4$) | [5], [2] |
| **Basophil** | 14-16 | Round, pleomorphic in tissue | Unknown | Bilobed | 1.4 (1.35-1.42) | $0.82*10^4$ (STD=$0.135*10^4$) | [3], [2] |
| **Macrophage** | 20-30 | Large, irregular, triangular | Unknown | Single-lobed, round centered | 1.384 ± 0.015 | Not Known | [6] |
| **Langerhans Cell** | 15-25 | Dendritic | Unknown | Indented "coffee bean" | Not Known | Not Known | [7] |

**Table S1. Distinct optical and cellular properties enable visualization and morphological distinction between major immune cell classes such as dendritic cells, macrophages and lymphocytes.**

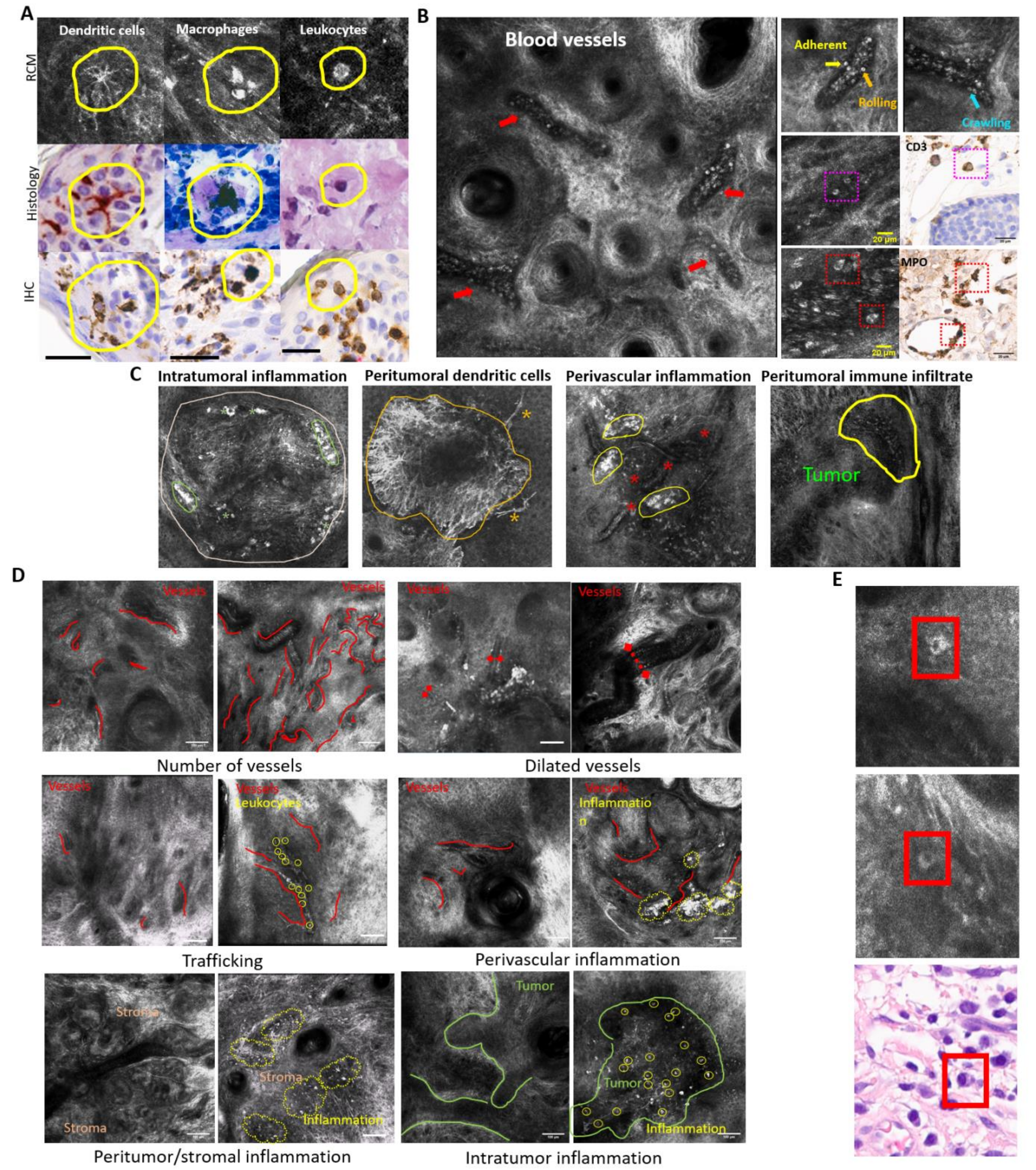


**Figure S1. Tumor-immune microenvironment and vasculature visualized and evaluated on RCM.**

**A.** Dendritic cells, macrophages and lymphocytes identified on RCM confirmed on histology and immunohistochemistry (CD1, CD68 and CD3, wall) in targeted biopsies. **B.** RCM images with blood vessels (red arrows) and trafficking of leukocytes on the vessel edges and individual leukocyte trafficking steps: adhering, rolling and crawling. Trafficking of lymphocytes and neutrophils confirmed using CD3 and Myeloperoxidase IHC, respectively. **C.** Intratumoral (green asterisk), and perivascular (yellow areas, outside blood vessels marked with red asterisk) peritumoral (orange asterisk, dendritic cells) distribution of inflammatory cells along with an immune infiltrate (yellow polygon) surrounding tumor seen on RCM. **D.** Representative example of features assessed during manual evaluation. For each feature, images with no/minimal feature presence and scored either 0 or 1 are shown on the left, while images with high density of features scored as 2 or 3 shown on the right (Red curve= blood vessels, diameter of vessels indicated by length of lines between diamonds; trafficking leukocytes encircled in yellow; inflamed area adjacent to vessels or in stroma marked by yellow lines; intratumor immune cells encircled in yellow) **E.** Round to elliptical, large (15-20 micron) immune cells with an eccentric nucleus and bright cytoplasm in dermis, often adjacent to blood vessels (perivascularly) resembling plasma cells on histopathology (red boxes) with an eccentric nucleus, perinuclear hoff and cytoplasm, seen next to a blood vessel. RCM: reflectance confocal microscopy; CD: cluster of differentiation; IHC: immunohistochemistry.

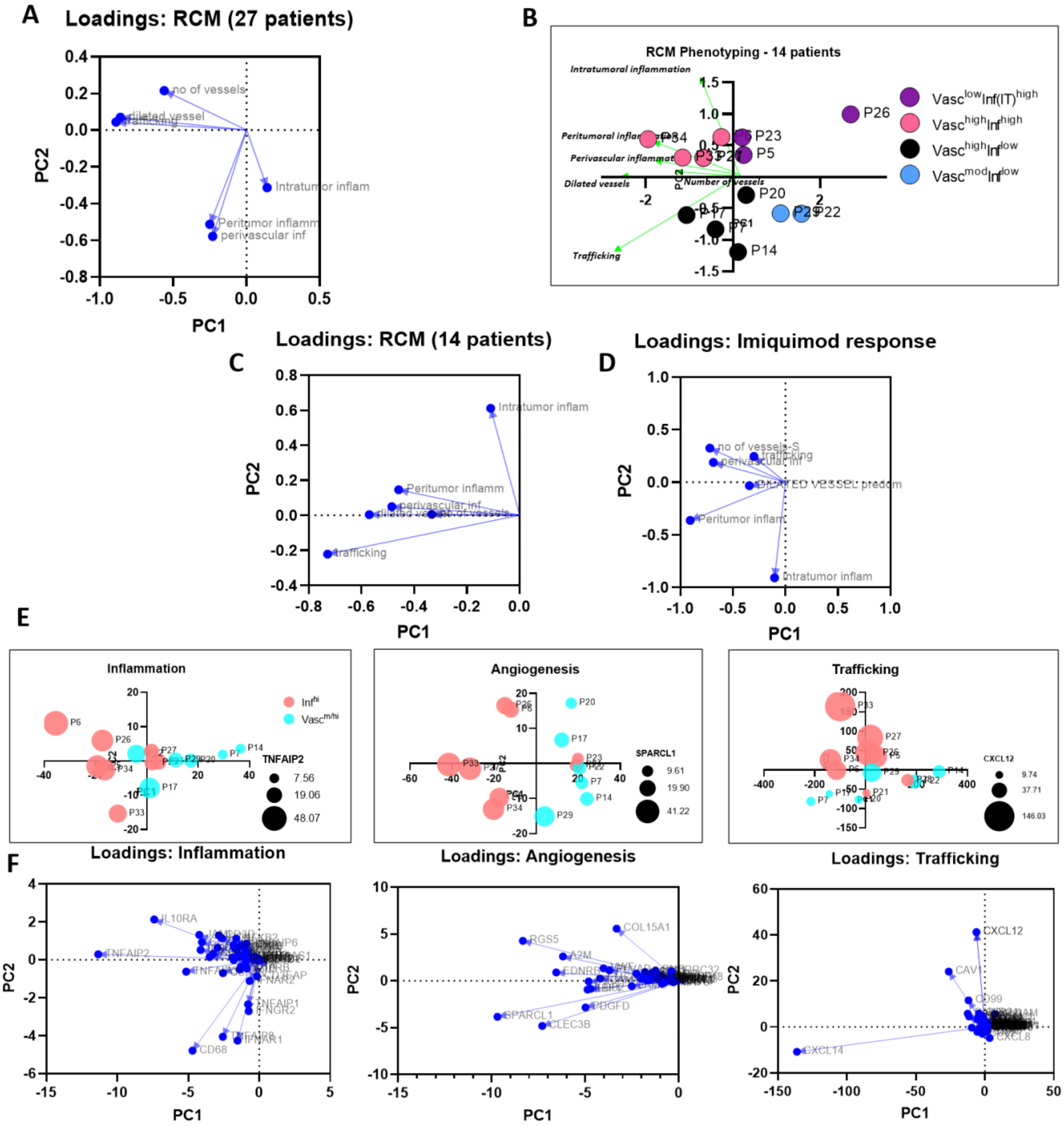


**Figure S2. PCA and PC loadings for RCM and gene expression analysis.**

**A.** PC loadings for PCA on total BCC patients. **B.** PCA on 14 BCC patients analyzed on gene expression analysis. **C.** PC loadings for PCA on BCC patients with gene expression. **D.** PC loadings for PCA on BCC patients undergoing imiquimod treatment. **E.** Unsupervised analysis (PCA) for phenotypes in inflammation, angiogenesis[8] and trafficking gene signatures demonstrate similar clustering as RCM phenotyping driven mainly by TNFAIP2, SPARCL1 and CXCL12, respectively inflammation, angiogenesis and trafficking genes. **F**. PC loadings for PCA on inflammation, angiogenic signatures and trafficking genes. PCA: principal component analysis; PC: principal components; RCM: reflectance confocal microscopy; BCC: basal cell carcinoma.

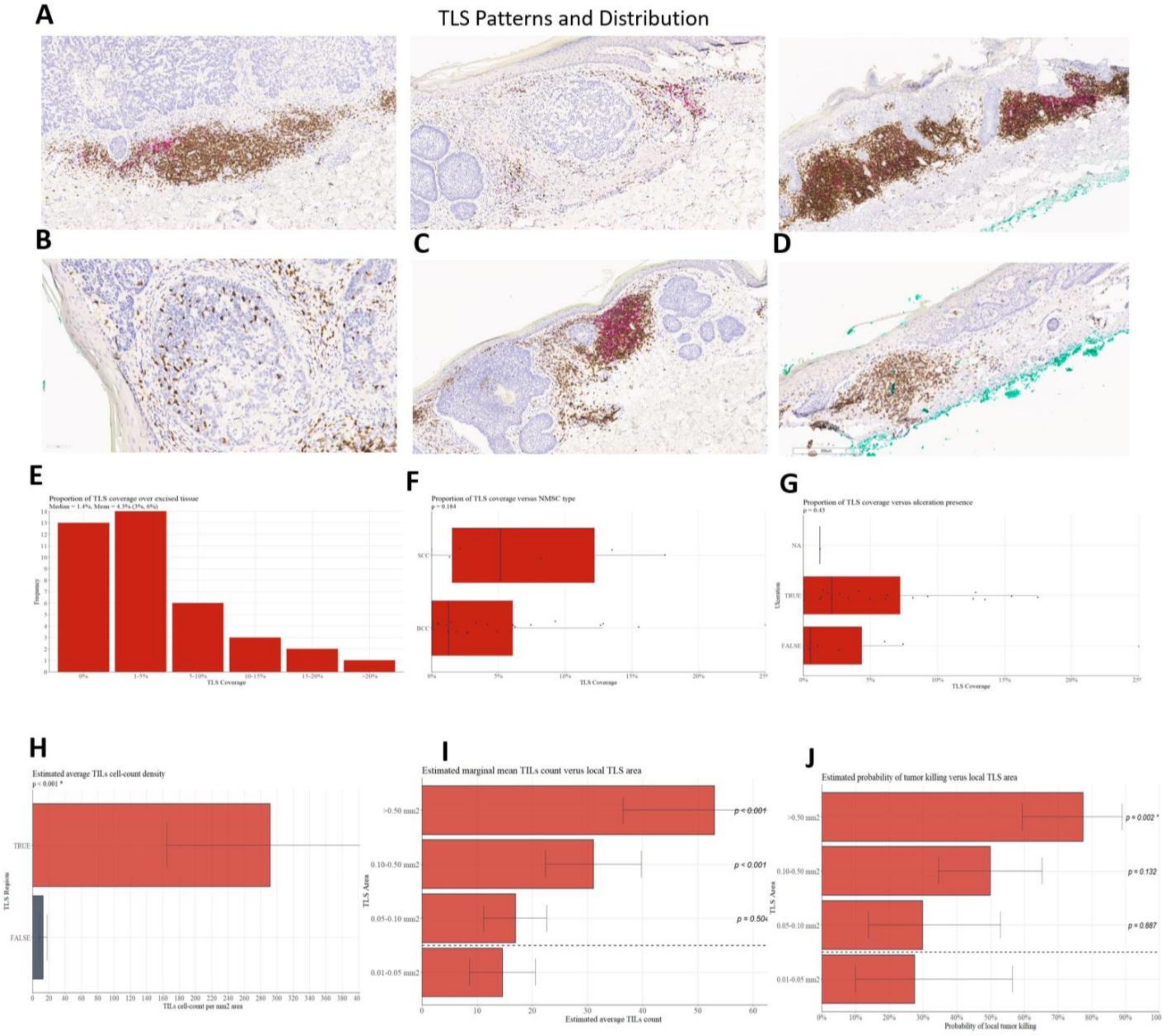


**Fig S3. Tertiary lymphoid structures abound in non-melanoma skin cancers and potentially represent anti-tumor immune response manifesting as higher TILs and tumor killing**

**A.** Representative images from $CD3^+/CD20^+$ staining of non-melanoma skin cancers (BCC and SCC) demonstrates differences in size, patterns and frequency of TLS in different patients. In addition to quantification of $CD3^+$, $CD20^+$ and total positive areas using software-based approaches, manual evaluation for number and size of TLS, local TILs and tumor killing were noted for a more spatial and context-dependent analysis. **B.** Representative image highlighting the high number of tumor-infiltrating $CD3^+$ cells inside BCC tumor nest adjacent to TLS (bottom). **C.** Large TLS surrounding remains of a tumor as confirmed by H&E. **D.** Less TILs and no tumor killing was observed in non-TLS patients and regions. **E.** Relative distribution of TLS in 40 BCC and SCC patients suggests a mean area coverage of 4.3% (CI: 3-6%). **F.** Higher TLS coverage in SCC as compared to BCC (p=0.18, ns). **G.** TLS presence corresponded with ulceration in most cases (p=0.43, ns). **H.** TLS-positive areas demonstrated higher TILs as compared to TLS-negative areas (p<0.001). **I.** Higher TLS coverage (>0.1 mm2) correlated with higher TILs (p<0.001). **J.** Higher TLS area coverage was associated with a statistically higher probability of tumor killing. TLS: tertiary lymphoid structures; TILs: tumor infiltrating lymphocytes, BCC: basal cell carcinoma: SCC: squamous cell carcinoma; CD: cluster of differentiation; CI: confidence interval.

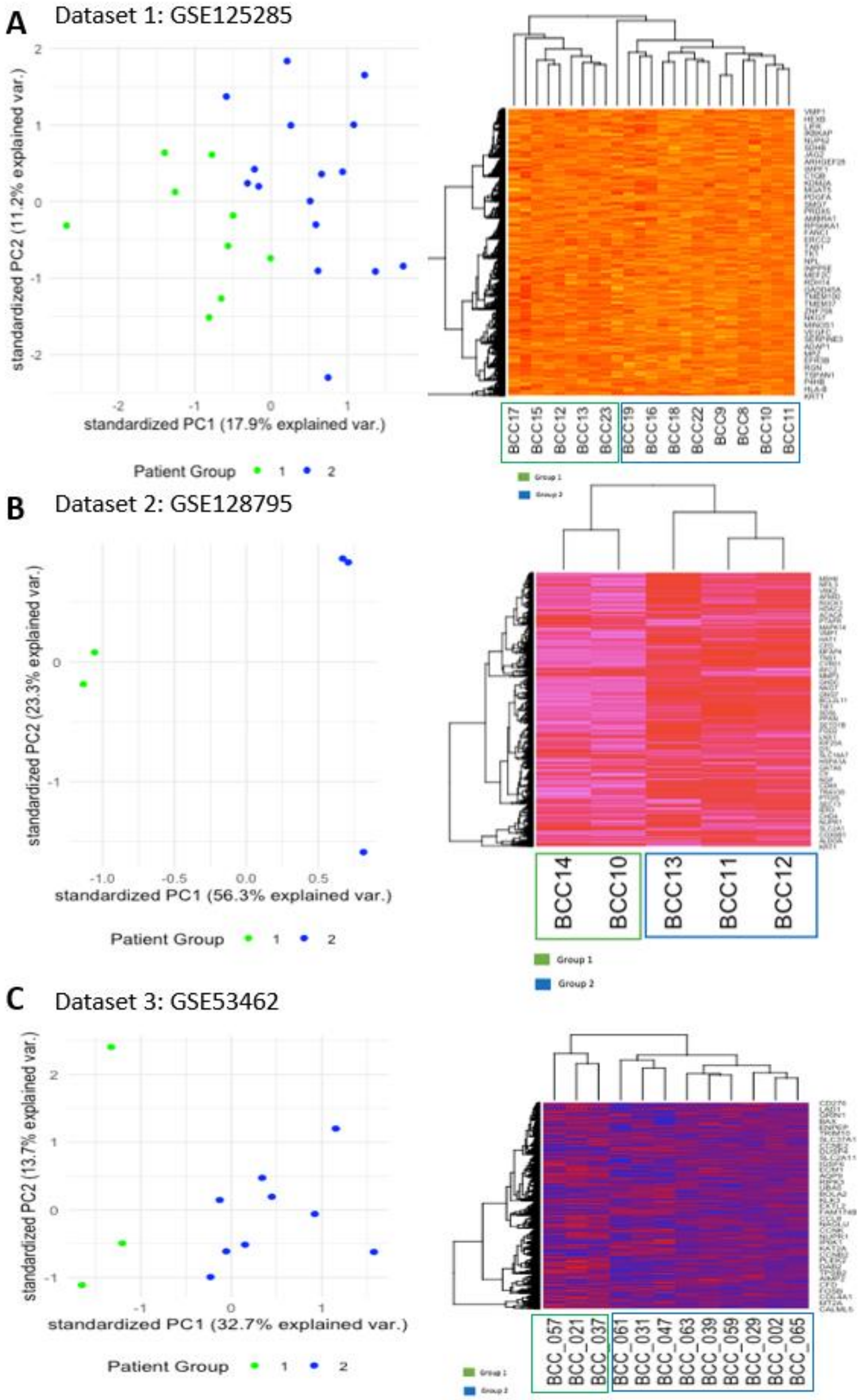


**Fig S4. Gene expression analysis on 3 distinct GEO datasets reveal presence of distinct phenotypes.**

**A.** PCA on GEO dataset GSE125285 Wan et al reveals presence of at least 2 phenotypes (green and blue). Unsupervised HCA also result in two phenotypes. GO on the DGEA demonstrate the following enriched (FDR < 0.01) terms: immune system process, cell surface receptor signaling pathway, lymphocyte activation, T cell activation, positive regulation of response to stimulus, leukocyte activation, positive regulation of immune system process, regulation of immune system process, and cell activation. **B.** PCA on GEO dataset GSE128795 Sand et al also highlights distinct immune phenotypes. Top GO Terms enriched (FDR <-0.05): leukocyte homeostasis, regulation of lymphocyte activation, cell adhesion, and biological adhesion. **C.** PCA on GEO dataset GSE53462 (Jee et al) also again reveals presence of at least 2 phenotypes (green and blue). Unsupervised HCA of 2 groups also results in two phenotypes. GO on the DGEA ( $|log2FC| > 0.3785$ and $FDR < 0.05$) demonstrate the following enriched ($FDR < 0.01$) GO terms: immune cell processes, antibacterial peptide production, antibacterial humoral response, antimicrobial humoral immune response mediated by antimicrobial peptide, extracellular matrix disassembly, and antimicrobial humoral response, perhaps suggestive of tertiary lymphoid structure and leukocyte trafficking heterogeneity in these patient clusters. *Only the subset of BCCs that did not display a "normal-like" or "SCC-like" gene expression pattern were kept in the analysis. PCA: principal component analysis (PCA); HCA: hierarchical cluster analysis; GEO: Gene expression omnibus; GO: gene ontology; FDR: false discovery rate; BCC: basal cell carcinoma; SCC: squamous cell carcinoma

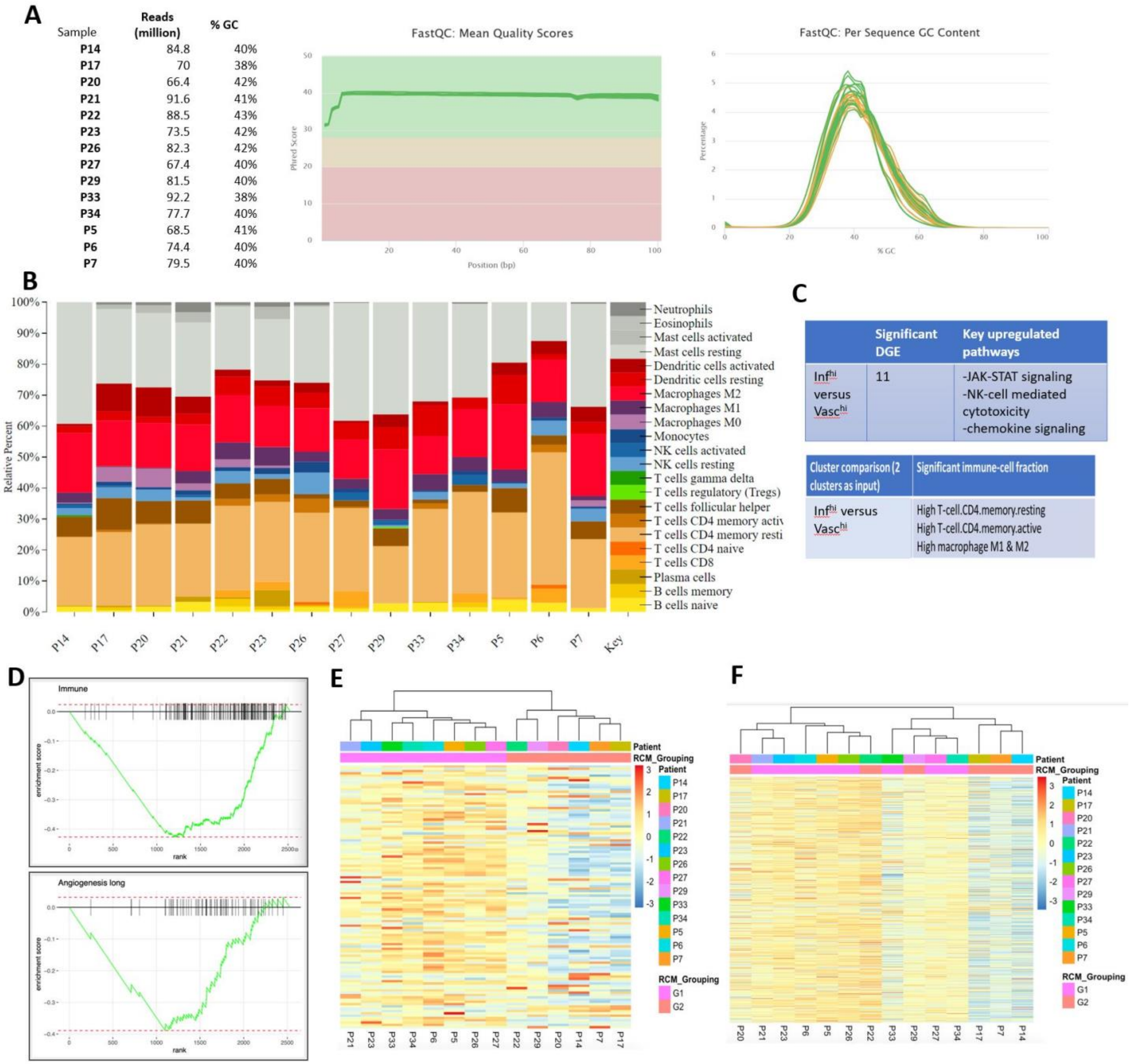


**Fig S5. Gene expression analysis using bulk RNA-seq demonstrates heterogeneity in gene expression**

**A.** Overall QC analysis suggests high quality and reliable transcriptomic data. **B.** CIBERSORT indicates differences in relative distributions of immune cell types in patients. **C.** DGEA on CIBERSORT output indicate upregulation of JAK-STAT, chemokine signaling and NK-mediated cell cytotoxicity in $Inf^{hi.}$ Statistically significant immune cells differences included CD4 memory resting and memory active cells, M1 and M2 macrophages**. D.** GSEA on $Vasc^{med/hi}$ and $Inf^{hi}$ demonstrates higher immune and angiogenesis signatures in $Inf^{hi}$. **E.** HCA on 114 genes differentially expressed genes. **F.** HCA on Nanostring Pancancer Immune Panel indicates presence of four phenotypes.

QC: quality control; DGEA: differential gene expression analysis; JAK-STAT: janus associated kinase- signal transducer and activator of transcription proteins; NK: natural killer; CD: cluster of differentiation; GSEA: Gene set enrichment analysis; HCA: hierarchical cluster analysis.

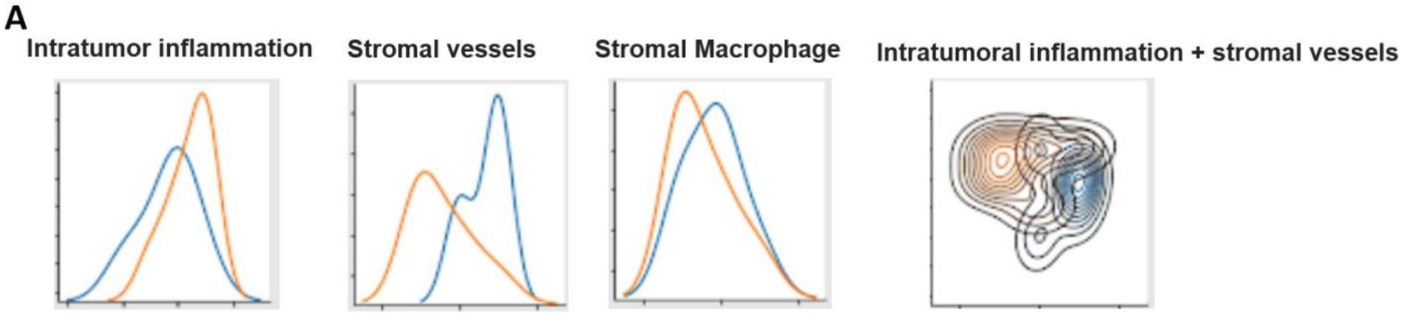


B

```
SPECIFICITY CRITERION
[2, 0, 3, 9, 10, 13, 16, 14, 5, 6, 4, 7, 18, 8, 17, 1, 15, 11, 12, 19]
[0.85714286 0.85714286 0.85714286 0.85714286 0.85714286 0.85714286
 0.85714286 0.85714286 0.71428571 0.71428571 0.71428571 0.71428571
 0.71428571 0.71428571 0.71428571 0.71428571 0.71428571 0.71428571
 0.71428571 0.71428571]
[0.66666667 0.66666667 0.66666667 0.66666667 0.66666667 0.66666667
 0.66666667 0.66666667 0.66666667 0.66666667 0.66666667 0.66666667
 0.66666667 0.66666667 0.66666667 0.66666667 0.66666667 0.5
 0.5        0.5       ]
[-15.06238189 -15.06238189 -15.06238189 -15.06238189 -15.06238189
 -15.06238189 -15.06238189 -15.06238189 -11.32251495 -11.32251495
 -11.32251495 -11.32251495 -11.32251495 -11.32251495 -11.32251495
 -11.32251495 -11.32251495  -8.42164879  -8.42164879  -8.42164879]

AIC CRITERION
[2, 5, 0, 4, 6, 7, 11, 13, 1, 16, 18, 8, 19, 3, 12, 14, 10, 15, 9, 17]
[0.85714286 0.71428571 0.71428571 0.71428571 0.71428571 0.71428571
 0.71428571 0.71428571 0.71428571 0.71428571 0.71428571 0.71428571
 0.71428571 0.57142857 0.57142857 0.57142857 0.57142857 0.57142857
 0.57142857 0.71428571]
[0.66666667 0.83333333 0.83333333 0.83333333 0.83333333 0.83333333
 0.83333333 0.83333333 0.83333333 0.83333333 0.83333333 0.83333333
 0.83333333 0.66666667 0.66666667 0.66666667 0.5        0.5
 0.33333333 0.5       ]
[-15.06238189 -15.06238189 -15.06238189 -15.06238189 -15.06238189
 -15.06238189 -15.06238189 -15.06238189 -15.06238189 -15.06238189
 -15.06238189 -15.06238189 -15.06238189  -8.42164879  -8.42164879
  -8.42164879  -6.05146855  -6.05146855  -4.04750971  -8.42164879]
```

**Figure S6. Modeling of imiquimod response highlights importance of stromal features in improving predictive power of linear regression models. A.** Linear separability curves for intratumor inflammation, stromal vessels and stromal macrophages (vessels and macrophages were predicted as priority labels in AIC and specificity prioritizing models) and combined plot for stromal vessels + intratumoral inflammation shows maximum separability (orange-responders, blue-non-responders). **B.** Models based on prioritizing Specificity or AIC as criteria for selecting features associated with response in linear regression models. For specificity, sequential modeling using 13 factors gave optimum performance while for AIC models the best performance was seen at 8 features. AIC: Akaike Information Criterion.

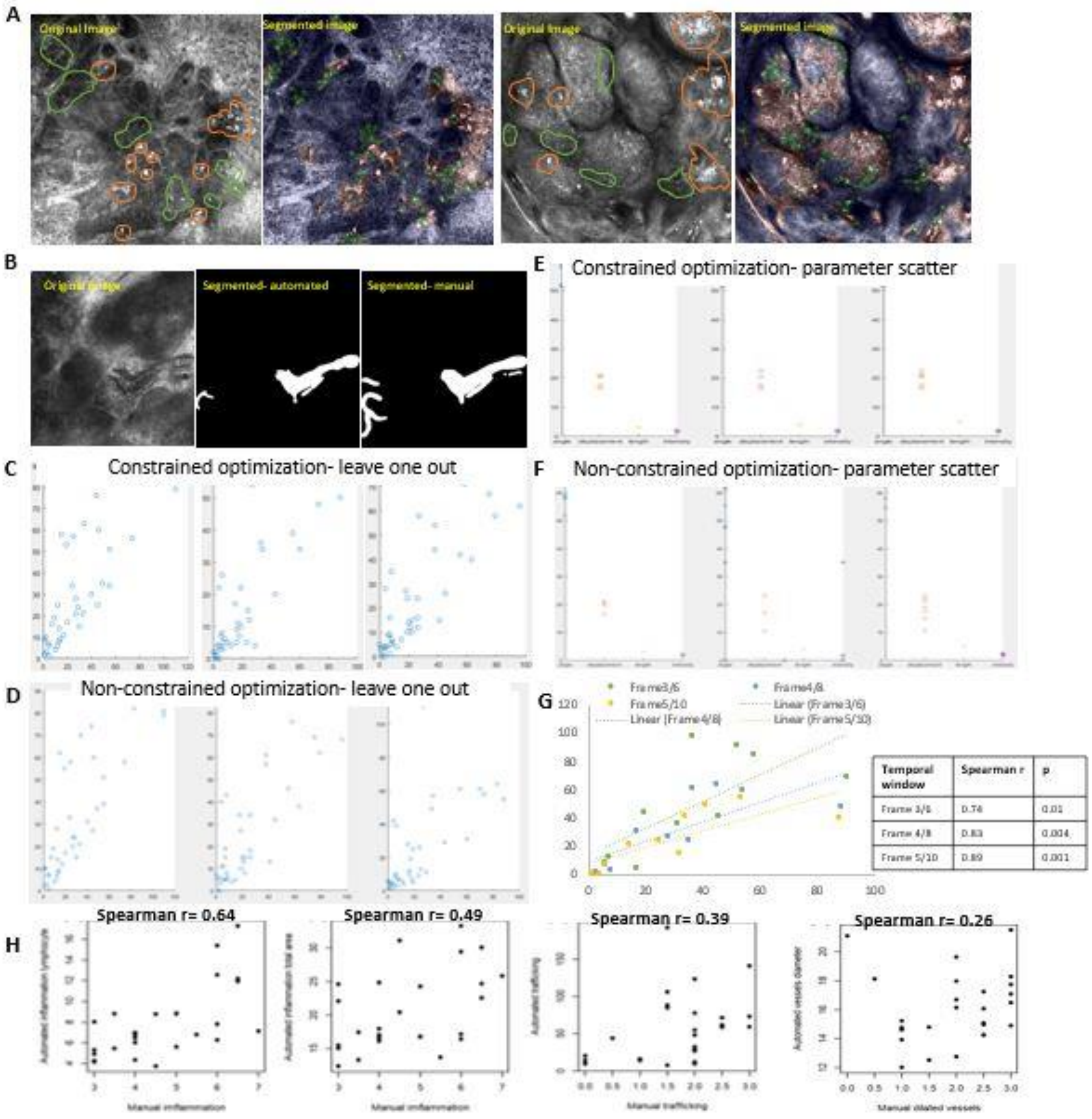


**Figure S7. Automated quantification of TiME and vasculature features on RCM images**

**A.** Exemplar RCM images (original and segmented image) showing segmentation of round-ellipsoid leukocytes (green), dendritic cells and macrophages (orange) using a UNet CNN model. **B.** Exemplar RCM video frames (original, manually segmented, automated segmented) showing segmentation of blood vessels from single field-of-view RCM videos. **C-F.** Summary of optimization for leukocyte trafficking done using constrained and non-constrained optimization of parameters: leave-one out and parameter scatter. **G.** High Spearman correlation (0.74-0.89) observed for average manual reader count and automated counts for all 3 temporal windows. H. Spearman correlation between automated quantification and manual reading. RCM: reflectance confocal microscopy; CNN: convolutional neural network.

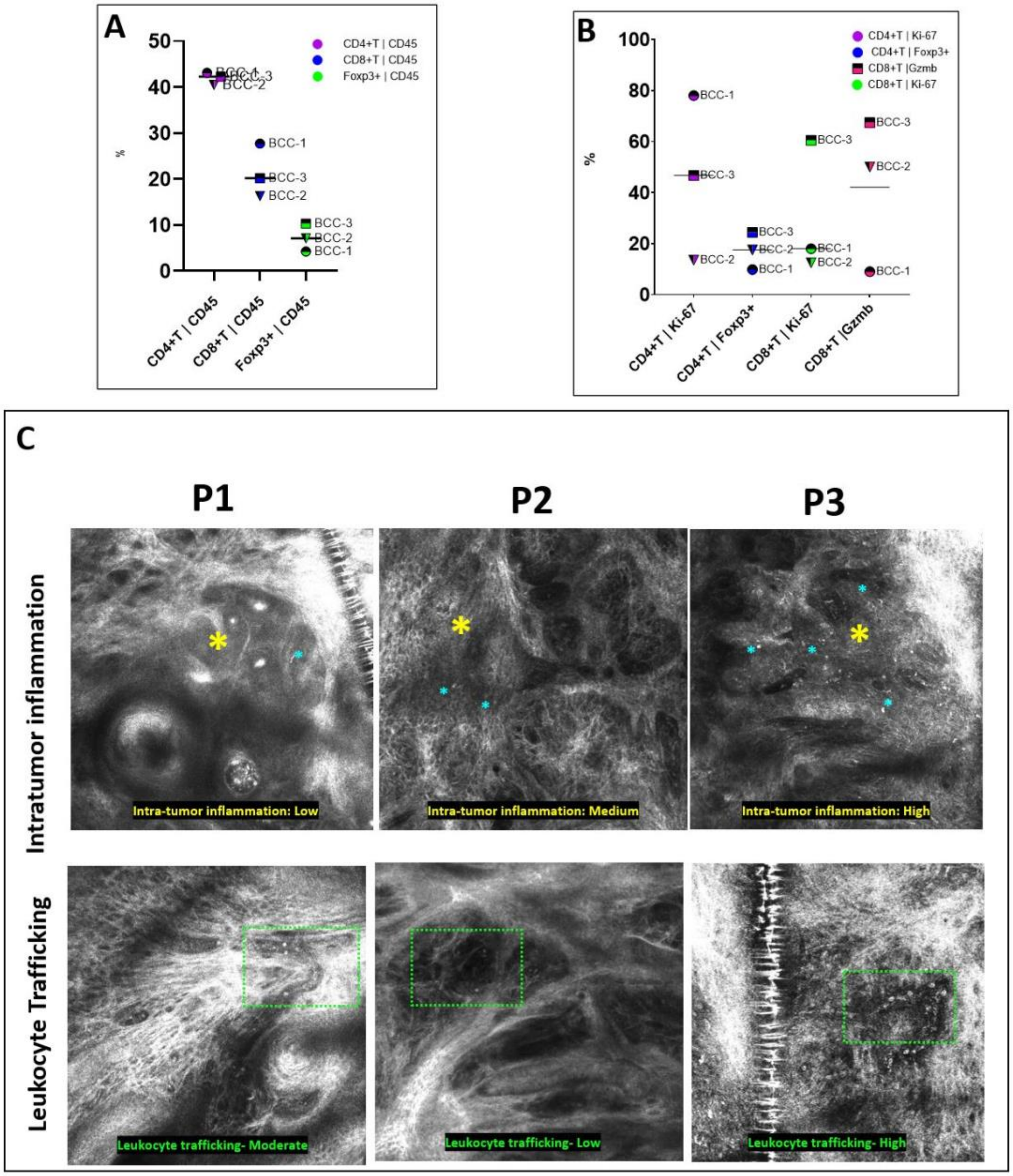


**Fig S8. Flow-based immunophenotyping for understanding T-cell distribution in red and blue RCM phenotypes.** **A.** Relative distribution of T-cell subsets- CD4, CD8 and Foxp3 in 3 BCC patients. **B.** Relative distribution of proliferating, activated or regulatory T cells. **C.** Representative RCM images from BCC1, 2 and 3 patients highlighting the differences in intratumoral inflammation, trafficking and vasculature between the Vasc$^{hi}$Inf$^{hi}$ (BCC-1) and Vasc$^{hi}$Inf$^{lo}$ (BCC-3) patient. BCC: basal cell carcinoma; RCM: reflectance confocal microscopy

**Supplementary videos:**

**Video 1:** Blood flow and leukocyte trafficking (within red border) in real-time within human dermis

**Video 2:** Rolling leukocytes (red arrowhead) in real-time within a blood vessel

**Video 3:** Crawling leukocyte (red arrowhead) in real-time within a blood vessel
